# Neural networks built from enzymatic reactions can operate as linear and nonlinear classifiers

**DOI:** 10.1101/2024.03.23.586372

**Authors:** Christian Cuba Samaniego, Emily Wallace, Franco Blanchini, Elisa Franco, Giulia Giordano

## Abstract

The engineering of molecular programs capable of processing patterns of multi-input biomarkers holds great potential in applications ranging from in vitro diagnostics (e.g., viral detection, including COVID-19) to therapeutic interventions (e.g., discriminating cancer cells from normal cells). For this reason, mechanisms to design molecular networks for pattern recognition are highly sought after. In this work, we explore how enzymatic networks can be used for both linear and nonlinear classification tasks. By leveraging steady-state analysis and showing global stability, we demonstrate that these networks can function as molecular perceptrons, fundamental units of artificial neural networks—capable of processing multiple inputs associated with positive and negative weights to achieve linear classification. Furthermore, by composing orthogonal enzymatic reactions, we show that multi-layer networks can be constructed to achieve nonlinear classification.

## I. Introduction

Cellular decision-making encompasses various endogenous processes that determine the fate of cells, whether they should replicate, differentiate, or undergo cell death. For example, apoptosis describes processes associated with programmed cell death to eliminate cells that are classified as abnormal or unnecessary cells. In embryonic development, individual cells process multiple environmental cues to decide whether they should differentiate, migrate, or decay. We are only beginning to understand the underlying mechanisms and factors influencing these crucial cellular decisions, which depend on a multitude of factors. Recent work has put forward the intriguing hypothesis that cellular decision making may present similarities with classification tasks performed by Artificial Neural Networks [1]–[4].

Theoretical work has illustrated that biochemical networks including sequestration or phosphorylation reactions can approximate ideal Artificial Neural Networks (ANNs) for linear and nonlinear classifiers [3], [4]. Experimental, bottom-up implementations of classifiers using biological molecules have been recently proposed in vitro, based on DNA strand displacement [5], and using the enzymatic PEN toolbox, leading to the demonstration of nonlinear classifiers [6]. In bacteria, synthetic genetic circuits based on transcription factors can generate diverse patterns [1], while metabolic networks can implement linear classifiers [7]. In mammalian cells, sequestration-based and and phosphorylation-based nonlinear classifiers have been demonstrated [8], [9]; further, protein-based linear classifiers based on the “winner-takes-all” principle have also been built [10].

Here, we examine a new set of biomolecular programs based on enzymatic association and degradation to build a biomolecular classifier. We take inspiration from the architecture of classifiers in ANNs, which are based on the interconnection of simple fundamental units known as “perceptrons”. A molecular perceptron requires only two core species, an enzyme and its substrate. The species producing enzyme and substrate serve as inputs to the perceptron. Through a rigorous steady-state analysis, we establish that our perceptrons operate robustly and generate classifiers with a linear *decision boundary* (namely, the hyper-surface that separates data points belonging to different classification labels), as long as enzyme-substrate association is sufficiently fast. Through computational simulations we then illustrate how the interconnection of multiple perceptrons with linear decision boundaries produces nonlinear classifiers. The simple enzymatic reactions we adopt expand the repertoire of available systems that can generate classifiers.

## II. A Molecular Perceptron

A perceptron is the fundamental unit of ANNs. Perceptrons process a weighted sum of their inputs, which is fed into activator functions, for example ReLU (Rectified Linear Unit) activation functions. In this section, we propose an enzymatic reaction that asymptotically can behave as a molecular perceptron. Although concentrations and rate constants are always positive, the network we propose can process multiple inputs associated with positive and negative weights. We provide a thorough analysis of the perceptron module dynamics, and in particular we show that the trajectories are bounded, and that the system admits an equilibrium that is structurally globally asymptotically stable.

### A. Chemical reactions and ODE model

In the following we denote chemical species with uppercase letters (e.g. *X*) and their concentrations with the corresponding lowercase letters (*x*). An enzymatic perceptron (Fig. 1A) consists of two species, *Y* (substrate) and *Z* (enzyme). Free (unbound) enzyme *Z* binds to the substrate *Y*, forming the enzyme-substrate complex *C*, at rate constant *γ*. This complex *C* catalyzes the degradation of *Y*, while *Z* dissociates from the complex, releasing free enzyme. The catalytic step (and release of *Z*) occurs at rate constant *θ*. Two inputs *X*_1_ and *X*_2_ produce respectively the species *Y* and *Z* at rate constants *w*_1_ and *w*_2_. We assume that all the species *Y, Z* and *C* dilute/degrade with rate constant *δ* as shown in Fig. 1A. Overall, our proposed enzymatic perceptron module is described by the following chemical reactions:

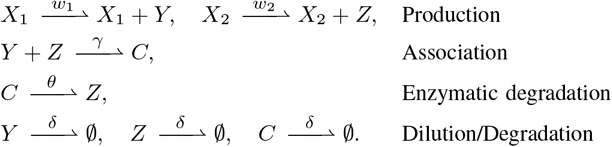

**Fig. 1.**
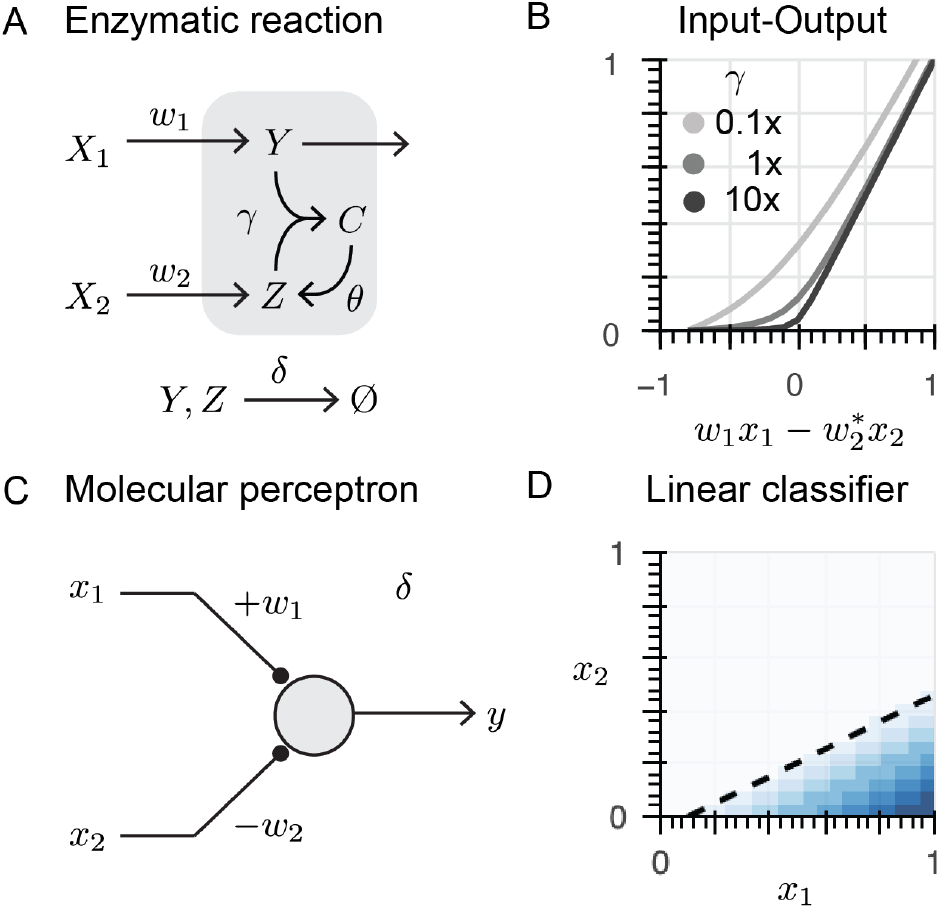
A: Reaction network describing the enzymatic perceptron. B: The input-output curve for different values of *γ* computes a ReLU activation function; 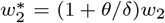. C: Schematic of the general architecture of a perceptron. D: At steady-state, the enzymatic perceptron realizes a linear classifier with a decision boundary corresponding to the black dashed line.

The association between enzyme and substrate is a second order reaction akin to molecular association [3], [11]–[13]. We will discuss the key role played by the related reaction rate parameter *γ* in guaranteeing desirable properties of this network.

Based on the law of mass action, we can write the corresponding system of Ordinary Differential Equations (ODEs) to describe the time evolution of the species concentrations:

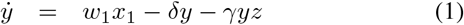

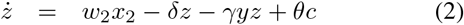

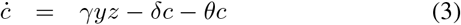

We now examine the stability properties of this circuit.

### B. Steady-state analysis in a fast association regime

We will show that as long as the enzyme/substrate association constant *γ* is sufficiently large, the system (1)-(3) asymptotically operates as a perceptron. In other words, this system processes the weighted sum of input *x*_1_ (associated with a positive weight) and input *x*_2_ (associated with a negative weight) to generate a piecewise linear function that is equal to the difference of the weigthed sum, if the difference is non-negative, and zero otherwise. This operation is equivalent to a ReLU activation function.

Let us compute the equilibria of system (1)-(3). By setting equation (3) to zero, we have:

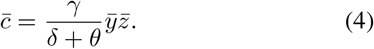

Then, 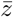 is obtained by setting both equations (1) and (2) to zero:

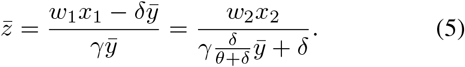

We thus obtain the second order polynomial

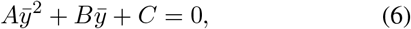

where

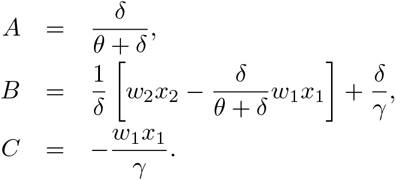

Since *C <* 0, equation (6) admits a single positive real solution, which is:

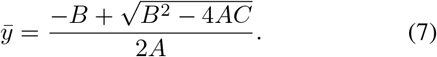

Relying on approximations that are valid in the fast association regime (i.e., *γ* arbitrarily large), we can consider the following approximated steady-state value for *y*:

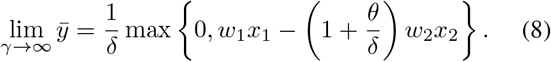

In fact, by defining *L* := lim_*γ→∞*_ − *B/A* and *M* := 4*w*_1_*x*_1_(*θ* + *δ*)*/δ*, we have

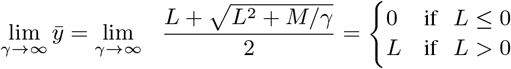

with

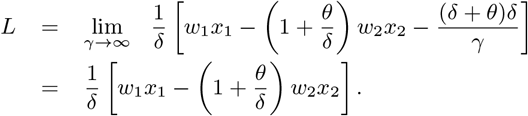

Equation (8) resembles the output of a ReLU activation function for two inputs: *x*_1_, with a positive weight +*w*_1_, and *x*_2_, with a negative weight 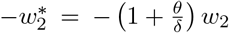, as shown in Fig. 1B. Decreasing the association rate constant *γ* makes the input-output response look like a soft-ReLU. When *θ* = 0, the enzyme-substrate association reaction (akin to molecular sequestration) is the core of the system. Therefore, we can abstract our enzymatic network as a molecular perceptron, as shown in Fig. 1 C.

The steady-state input-output map of system (1)-(3) can be computed by evaluating equation (7) when the inputs (*x*_1_ and *x*_2_) range from 0 to 1. In the heat plot shown in Fig. 1 D, the white region corresponds to an output close to zero, while the blue range corresponds to an output larger than zero. The black dashed line represent the *decision boundary* that separates the white and blue regions. This computed input-output map is equivalent to a linear classifier, because the decision boundary is a line dividing the input domain in two regions. The system output produces a response larger than zero only when the input combination (*x*_1_, *x*_1_) falls in the blue region.

We can also find a closed-form expression for the decision boundary (the black dashed line in Fig. 1 D). For this purpose, we rewrite equation (5):

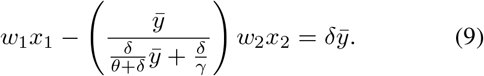

This equation describes the relationship between *x*_1_ and *x*_2_ when *y* is constant, as a function of the network reaction rate parameters. Fig. 1 D shows the decision boundary for 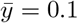. In the fast association regime (*γ→ ∞*), the decision boundary becomes:

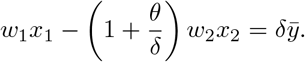

### C. Boundedness and equilibria

All the state variables of system (1)-(3) are bounded, for all possible nonnegative initial conditions, if the inputs *w*_1_*x*_1_ =: *α* and *w*_2_*x*_2_ =: *β* are constant and bounded. In fact, since the system is positive (each state variable has a nonnegative derivative when its value is zero), *y, z, c ≥* 0. Moreover,

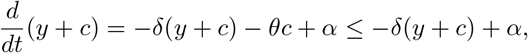

hence *y* + *c* is bounded, and similarly

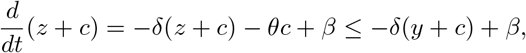

hence *z* + *c* is bounded. Therefore both *y* and *z* are bounded, because all the variables are nonnegative. Finally, from (3), *c* is also bounded.

Rewriting the dynamics of system (1)-(3), assuming constant inputs, with the new state variables *υ* = *y* + *c*, and *ζ* = *z* + *c* yields the ODE system

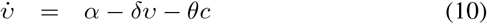

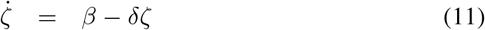

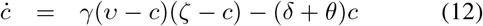

Since *y, z, c* are nonnegative variables, we have

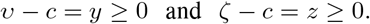

The system (10)-(12) has bounded solutions, for all possible nonnegative initial conditions with *c≤* min *{υ, ζ}* . In fact, *ζ* is bounded as it obeys to equation (11); then also *c* is bounded by *ζ*; and, from equation (10), also *υ* is bounded, because *c* is bounded.

Boundedness implies the existence of an equilibrium [14]– [16]. For system (1)-(3), the equilibrium has been computed in Section II-B. For system (10)-(12), the equilibrium can be computed as follows. First, from (11), 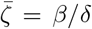. Then, by setting to zero equation (12), we obtain

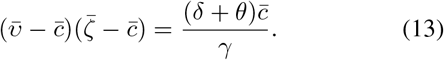

Since *c* is bounded by *ζ* regardless of the value of *γ*, for *γ→ ∞* the right-hand-side of equation (13) converges to 0, and hence the limit equation is (*υ− c*)(*ζ − c*) = 0, which means that either *υ − c* = 0 or *ζ − c* = 0. On the other hand, since *c* is upper bounded by both *υ* and *ζ*, for *γ→ ∞* the equilibrium becomes

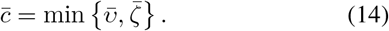

Now we can distinguish two cases.

First case: if 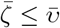, then 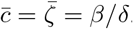.

Second case: if 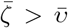, then 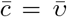. By setting to zero equation (10), we get

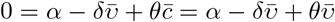

and hence 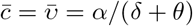.

Therefore, in the fast association regime (*γ → ∞*),

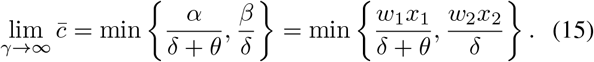

Then, the equilibrium value for *υ* in the fast association regime is

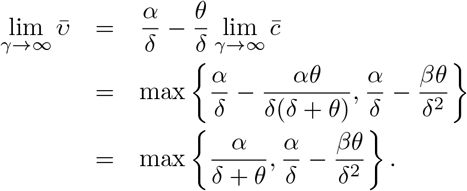

### D. Stability analysis

We now assess the stability of the equilibrium that we have computed. Since system (10)-(12) is obtained from system (1)-(3) through a linear coordinate transformation, the stability properties of the equilibrium are identical in the two cases.

We start by proving its local asymptotic stability, which holds structurally, i.e., regardless of parameter values [17].

#### Proposition 1.

*The equilibrium point of system* (10)-(12) *(equivalently, of system* (1)-(3)*) is structurally locally asymptotically stable*.

*Proof*. At steady-state, we have that 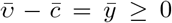 and 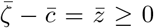, while 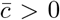. Therefore, we can prove that the equilibrium 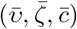 is structurally asymptotically stable [17], regardless of parameter values. Indeed the Jacobian of system (10)-(12) is

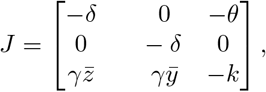

where 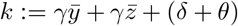. Its characteristic polynomial *ψ*(*s*) = det(*sI − J*) is Hurwitz, since it can be factored out as 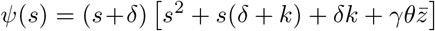, and therefore all its roots must have strictly negative real part. □

We can actually prove *global* stability of the equilibrium, again structurally [17]. Consider again the original representation, which we report here for convenience:

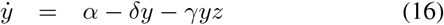

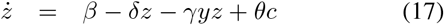

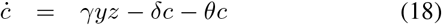

where *α* and *β* are constant positive inputs. We recall that the shifted variable

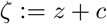

satisfies the differential equation

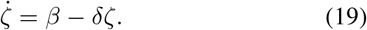

Hence, *ζ* (namely, *z* + *c*) exponentially converges to the variety

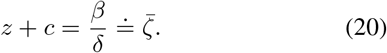

We can then prove convergence inside this variety. Consider the first two equations and write them as

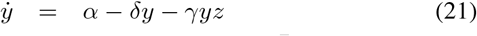

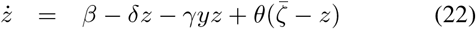

Denoting by 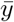 and 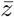 the equilibrium values corresponding to the given *α* and *β*, we now consider the shifted system in the variables 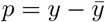 and 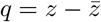:

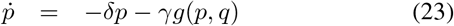

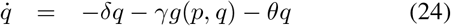

where we denoted 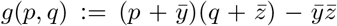. Since *g*(0, 0) = 0, the new system admits the equilibrium (*p, q*) = (0, 0).

To structurally prove global asymptotic stability, we apply the theory developed in [18], [19]. We write

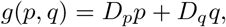

where *D*_*p*_(*p, q*) and *D*_*q*_(*p, q*) are positive functions; for instance,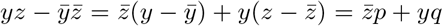. Then, the system can be written as

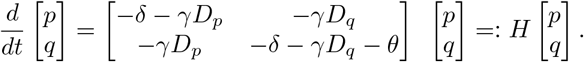

Since matrix *H* is column diagonally dominant and strictly nonsingular, it admits the 1-norm as a Lyapunov function. Therefore, the asymptotic stability of the equilibrium is global, and it is structural since it holds regardless of the values of the positive parameters.

We formalize these results in the following statement.

#### Proposition 2.

*For any initial condition and constant positive inputs, and for all values of the positive parameters, the state of system* (1)*-*(3) *(equivalently, of* (10)*-*(12)*) converges to the affine variety* (20). *Then, for any initial state in this variety, the system state structurally converges to the (unique) equilibrium. Therefore, the equilibrium is structurally globally asymptotically stable*.

## III. Multi-input Linear Classifier

In the previous sections we examined a perceptron that achieves linear classification by only processing two inputs *x*_1_ and *x*_2_. Here, through numerical simulations, we compute the decision boundary for perceptrons with three inputs, as shown in Fig. 2 A. The system now includes three inputs *X*_1_, *X*_2_, and *X*_3_, associated with positive weights *w*_1_, *w*_2_, and *w*_3_. Each of the inputs produces species *Y* . We introduce a *bias* input species *X*_0_, associated with a negative weight *w*_0_, which produces species *Z*. The association reaction between *Y* and *Z* is the same as in the two-input perceptron. All species dilute/degrade at the same rate constant *δ*. We list below all the chemical reactions networks, which reflect the abstract design in Fig. 2 A:

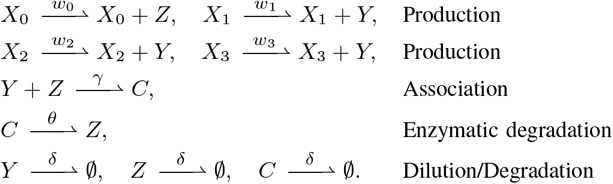

**Fig. 2.**
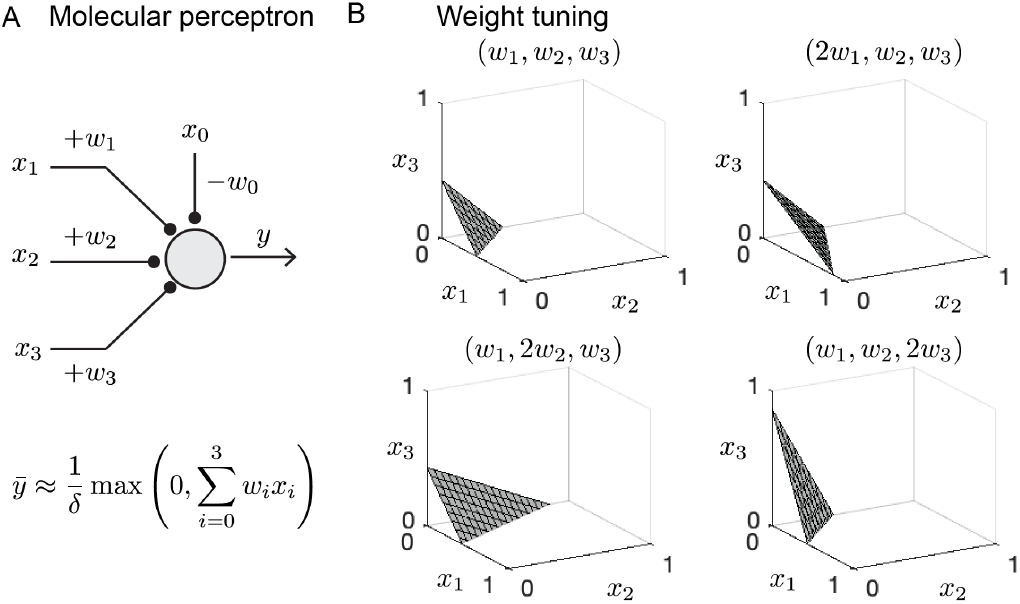
A: Schematic of a three-input perceptron. B: Simulations showing how the decision boundary depends on the input weights.

Next, using mass conservation, we can write the ODEs:

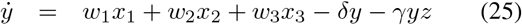

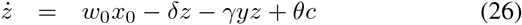

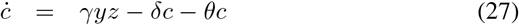

Following similar steps as in the perceptron case, we can compute the decision boundary at the steady-state, which for the multi-input linear classifier becomes

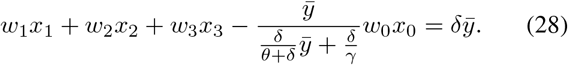

Equation (28) describes a plane in the space *x*_1_, *x*_2_, *x*_3_, assuming large association constant (*γ → ∞* ), as shown in Fig. 2B. When the combination of inputs is located in the right side of the plane, the output is larger than zero. In contrast, when the combination of input corresponds to a point below the plane, the output is zero. By changing the weights of the inputs, we can tune the decision boundary as shown in Fig. 2B.

## IV. Nonlinear Classifier

In this section, we report computational simulations that illustrate the behavior of multiple interconnected perceptrons. Through these interconnections, we show that we can create classifiers with nonlinear decision boundaries.

For this purpose, we design a two-layer classifier as shown in Fig. 3 A. The first layer consists of two enzymatic perceptrons, labeled 1 and 2 and represented as gray nodes in the perceptron network in Fig. 3 A and B. The first node process the inputs *x*_1_ and *x*_2_ with positive weight 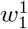 and a negative weight 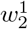 respectively; this node also processes the bias input *x*_0_ with a negative weight 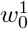. The second node takes in the inputs *x*_1_ and *x*_2_ with positive weights 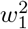 and 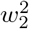, and it processes the bias input *x*_0_ with a negative weight 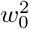. The second layer consists of a single node, represented in green in Fig. 3 A and B. So far, we have taken the substrate species *Y* as the output of the perceptron. Here, we take the complex *C* as the output, and show that we can produce a nonlinear decision boundary based on the outputs of the first layer. In this layer, we remove the enzymatic degradation step, but preserve the molecular association at the core of this node. In fact, from equation (15) we see that in the fast association regime, even when *θ* = 0, the complex output is the minimum of the two inputs, suitably scaled.

**Fig. 3.**
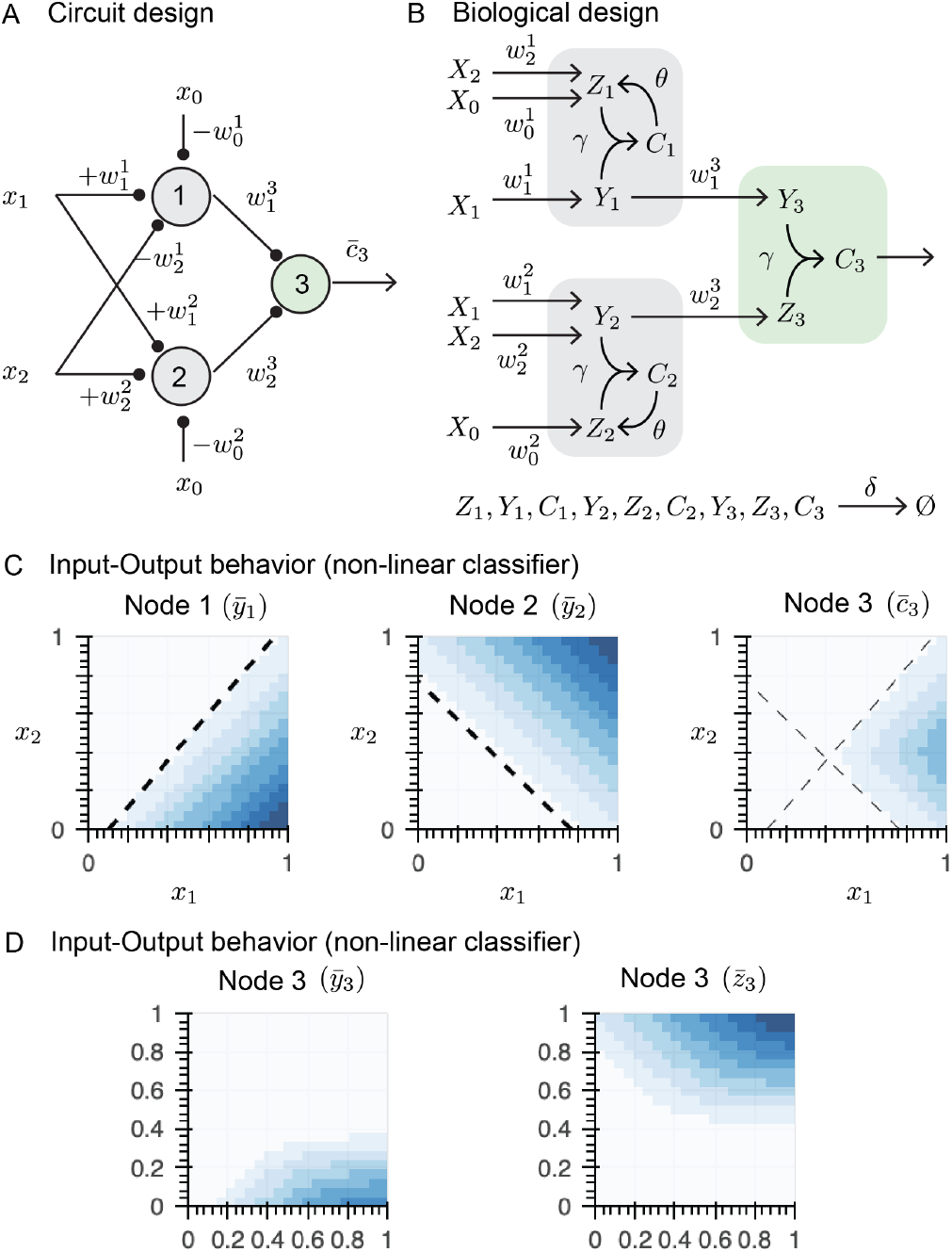
A: Schematic of a two-layer classifier. B: Reaction schematics corresponding to each node of the classifier. C: Computationally generated input-output map of each node. Node 3 shows a nonlinear decision boundary. D: Input-output behavior observed when the output of node 3 is either the substrate or the enzyme.

Using the reactions represented in the schematic in Fig. 3 B, we can derive the ODEs describing the dynamics of the first node:

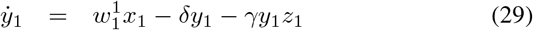

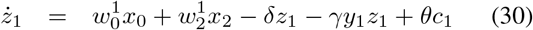

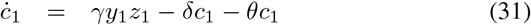

At steady-state, we can find the decision boundary, which is described by the linear equation:

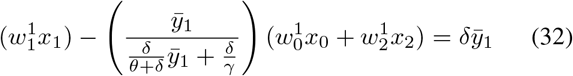

The input-output steady-state map of the first node is shown in Fig. 3 C, left. The decision boundary in Equation (32) is the dashed line separating the blue region from the white area. Similarly, given the schematic in Fig. 3 B we can find the ODEs describing the dynamics of node 2 in layer 1:

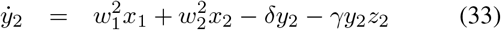

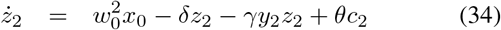

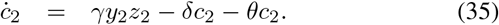

As we did before, we find the equation that describes the decision boundary for the node:

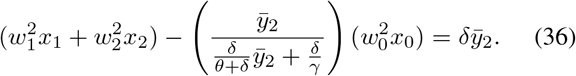

Fig. 3 C, middle, shows the steady-state input-output map of the second node. For both nodes 1 and 2, the computed decision boundaries separate the white region from the blue region. Input combinations associated with points in the plane above the decision boundary correspond to node outputs larger than zero. In contrast, input combinations associated with points below the decision boundary correspond to an output equal to zero.

Finally, we can write the equations corresponding to the second layer, which includes a single node that has the complex *c*_3_ as an output:

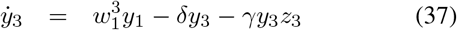

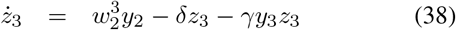

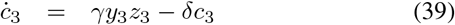

Resorting to the same derivations as in Section II-C, we find that, in the fast association regime, the output *c* computes the minimum of its inputs, suitably scaled, at steady-state. In particular, taking *θ* = 0, equation (15) leads to:

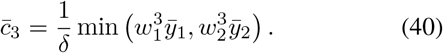

Now, by interconnecting the input-output maps of the first layer with the second layer, we can predict the overall output of the classifier. The output of node 3 is the scaled minimum of its two inputs. Therefore, when the output of either node 1 or node 2 is in the white region, the output *c*_3_ is also in the white region. The overall decision boundary for node 3 (i.e., the second layer) can be visualized by overlapping the decision boundaries of nodes 1 and 2 (i.e., the first layer), as shown in Fig. 3 C (right), which yields a nonlinear blue area. Thus, the combination of the linear decision boundaries from the first layer (node 1 and 2) sets the boundaries of the output of the node 3. For this reason, by tuning the individual linear decision boundaries of each node, we can tune the nonlinear decision boundary of the whole network. If the output of node 3 is either the substrate *y*_3_ or the enzyme *z*_3_, then the classifier retains a nonlinear decision boundary as shown in Fig. 3 D, however its shape can no longer be intuitively mapped to the boundaries of nodes 1 and 2.

**TABLE I.** Table 1. Parameters used in computational simulations.

| Parameter | Unit | Value | Figure |
| --- | --- | --- | --- |
| $\delta$ | $\text{h}^{-1}$ | 1 | all |
| $\theta$ | $\text{h}^{-1}$ | 1 | all |
| $\gamma$ | $\mu\text{M}^{-1}\text{h}^{-1}$ | 100 | all |
| $w_1 = w_2$ | $\text{h}^{-1}$ | 1 | Fig. 1 |
| $w_1 = w_2 = w_3$ | $\text{h}^{-1}$ | 1 | Fig. 2 |
| $w_0 x_0$ | $\mu\text{Mh}^{-1}$ | 0.6 | Fig. 2 |
| $w_1^1 = 2w_2^1$ | $\text{h}^{-1}$ | 1 | Fig. 3 |
| $w_0^1 x_0$ | $\mu\text{Mh}^{-1}$ | 0 | Fig. 3 |
| $w_1^2 = w_2^2$ | $\text{h}^{-1}$ | 1 | Fig. 3 |
| $w_0^2 x_0$ | $\mu\text{Mh}^{-1}$ | 0.4 | Fig. 3 |
| $w_1^3 = w_2^3$ | $\text{h}^{-1}$ | 1 | Fig. 3 |
| $w_0^3 x_0$ | $\mu\text{Mh}^{-1}$ | 0 | Fig. 3 |

## V. Discussion

We have demonstrated that suitably designed enzymatic reactions can asymptotically give rise to an ideal perceptron implementing a linear classifier, when the association rate is sufficiently large, and can process multiple inputs with both positive and negative weights. By composing multiple layers, we can realize nonlinear classifiers in a predictable, tunable and intuitive manner. This work expands the repertoire of sequestration- and phosphorylation-based Biomolecular Neural Networks that can realize complete ideal ANNs. In vitro strand displacement Neural Networks can only be used to design linear classifiers [5], and it is challenging to assemble multi-layer neural networks to get nonlinear classifiers or to translate such circuits into living systems. PEN neural networks are challenging to implement in living systems because of the non-ideal operating conditions they require [6]. Although transcriptional-based neural networks can realize nonlinear decision making [1], [20], they do not behave as ideal perceptrons, which makes it difficult to compose multiple layers and optimize computationally. Winner-takes-all networks [10] are limited to linear classification due to the multiple interconnections that arise from each implementation. These limitations raise the open question: what are the key design principles of molecular perceptrons?

The realization of molecular perceptrons based on various mechanisms – such as sequestration [3], phosphorylation [4], and enzymatic reactions (as in this work) – suggests that endogenous networks may be able to perform pattern recognition. However, the networks constructed so far lack the complexity of endogenous networks, which arises e.g. due to combinatorial binding [2], shared resources [21], and other cellular contexts. This complexity makes it challenging to evaluate how different endogenous networks are from an ideal ANN; this is a fascinating avenue for future work.

## Notes

### Competing Interest Statement

The authors have declared no competing interest.

